# ANTIMICROBIAL ACTIVITIES OF VERNONIA AMYGDALINA AGAINST BIOFILM-FORMING MICROBES

**DOI:** 10.1101/2025.05.22.655479

**Authors:** Ebiloma Samuel, Nkechi Chuks Nwachuckwu, Happy Uchendu Ndom, Hope Okereke

## Abstract

The increase threat of antimicrobial resistance, mainly among biofilm-forming pathogens, has prompted increased interest in plant-based alternatives with broad-spectrum antimicrobial properties. This study aimed to evaluate the antimicrobial activities of aqueous, ethanolic, and methanolic leaf extracts of *Vernonia amygdalina* against five biofilm-forming clinical isolates: *Staphylococcus aureus, Escherichia coli, Klebsiella pneumoniae, Pseudomonas aeruginosa*, and *Candida albicans*. Plant extracts were obtained through maceration using respective solvents and tested at concentrations ranging from 100 to 500 mg/mL. The antimicrobial potential of each extract was assessed using the agar well diffusion method to determine the zone of inhibition, followed by minimum inhibitory concentration (MIC) and minimum bactericidal/fungicidal concentration (MBC/MFC) assays. Among the three extract types, the methanolic extract consistently showed the highest antimicrobial activity, with inhibition zones ranging from 20.0 ± 0.38 mm to 24.0 ± 0.60 mm. Ethanolic extracts showed moderate inhibition, while aqueous extracts were least effective, particularly at lower concentrations. MIC values for methanolic extracts ranged from 31.25 mg/mL for *C. albicans* to 500 mg/mL for *S. aureus*, with corresponding MBC values as low as 31.25 mg/mL for *E. coli*. One-way ANOVA indicated a statistically significant difference (*p* < 0.05) in mean inhibition zones across extract types. Post hoc Tukey’s test confirmed the superiority of methanol as an extraction solvent. These results validate the broad-spectrum antimicrobial properties of *V. amygdalina* and support its potential development as a natural therapeutic agent against resistant, biofilm-forming microbial infections.

## 1. Introduction

The increase and the global threat of antimicrobial resistance has become a worldwide public health threat, compromising and altering the therapeutic effect of antimicrobial medications (world Health Organization [WHO], 2023).biofilm formation enhance the antimicrobial resistance, a microbial means of persistence that contribute to almost 80% of severe infections (Lebeaux et al., 2024). Biofilms are 20 to 1000 times more resistant to antimicrobial agents than their planktonic counterpart, because, biofilms are structured community of microbes covered in extracellular polymeric substances (Smarma et al., 2023; Pino et al., 2023).

Due to phytochemical diversity and long standing use in conventional medicine, medicinal plants are increasingly being investigated as root of new antimicrobial compounds (Negm et al., 2023). Tannins, terpenoids, flavonoids, alkaloids, phenolics are plants bioactive based compound that have shown impressive antimicrobial and antibiofilms activities (Elekhnawy et al., 2023; Obaod et al., 2023). These bioactive composites work through mechanisms including disruption of microbial membranes, inhibition of nucleic acid synthesis, and interference with quorum sensing (Zaman & Smith, 2025; Satria et al., 2023).

*Vernonia amygdalina* is rich in bioactive substances which contribute to pharmacological treatment of biofilm and microbial infection, it contain substances like flavonoids, phenolic acids, saponins and sesquiterpene (Ugboko et al., 2022; Adeleke et al., 2024). It is a perennial shrub known to be used as antimalarial, treatment of gastrointestinal disorders, microbial infections, and diabetes in sub-saharan Africa (Onohuean et al., 2023) the current research has shown the antimicrobial healing effect of *Vernonia amygdalina* leaf extracts against multidrug-resistant (MDR) pathogens such as *Escherichia coli, salmolla typhi*, and *Staphylococcus aureus* (Onohuean et al., 2023; Satria et al., 2023). Also, fermented extracts have shown improved antibacterial action, attributed to enhanced bioavailability of active compounds (Adeleke et al., 2024; Obaid et al., 2023). Methanolic and ethyl acetate extracts, in particular, have shown synergistic activity when combined with antibiotics like tetracycline and ciprofloxacin (Adeleke et al., 2024; Elekhnawy et al., 2023) beyond being an antibacteria agents *V. amygdalina* shown an antifungal activities, mainly against strong biofilm forming fungi such as *Candidal albicans* in immunocompromised patients (Negm et al., 2023). The biofilm inhibition potential of *V*.*amygdalina* is also reinforced by silico docking studies, exposing the relationships with microbial adhesion and quorum sensing pathways (Pinto et al., 2023; Sharma et al., 2023).

Despite these promising findings, studies focused on the antibiofilm activities of *V. amygdalina* against clinical isolates remain limited. Most existing works assess crude antimicrobial activity without comparing the efficacy of different extract types (aqueous, ethanolic, and methanolic) or determining minimum inhibitory concentrations (MIC) and bactericidal endpoints (MBC/MFC) (Ugboko et al., 2022; Satria et al., 2023).

Given the threat posed by biofilm-related infections, particularly in immunocompromised and hospitalized patients, it is imperative to explore affordable and accessible plant-based solutions. This study aims to evaluate the antimicrobial activities of *V. amygdalina* extracts against biofilm-forming microbes isolated from wound and cystic fibrosis patients, with emphasis on extract type, potency, and microbial specificity (WHO, 2023; Zaman & Smith, 2025; Adeleke et al., 2024).

## Findings

The antimicrobial activities of aqueous, ethanolic, and methanolic leaf extracts pf Vernonia Amygdalina against 5 biofilm forming microbes isolated from wounds, microbes like: *Staphylococcus aureus, Escherichia coli, Klebsiella Pneumoniae, Candida albicans, Pseudomonas aeruginosa*. The result presented below (Table 1) is the result obtained from well diffusion, minimum inhibitory concentration (MIC) and minimum bactericidal/fungicidal concentration (MBC/MFC).

**Table 1.**
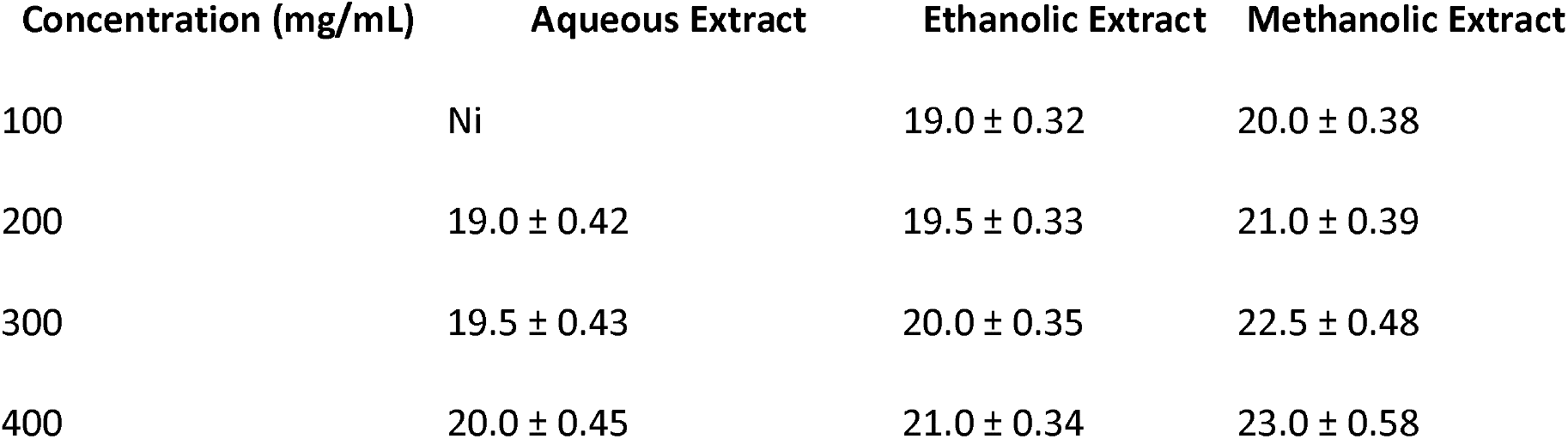
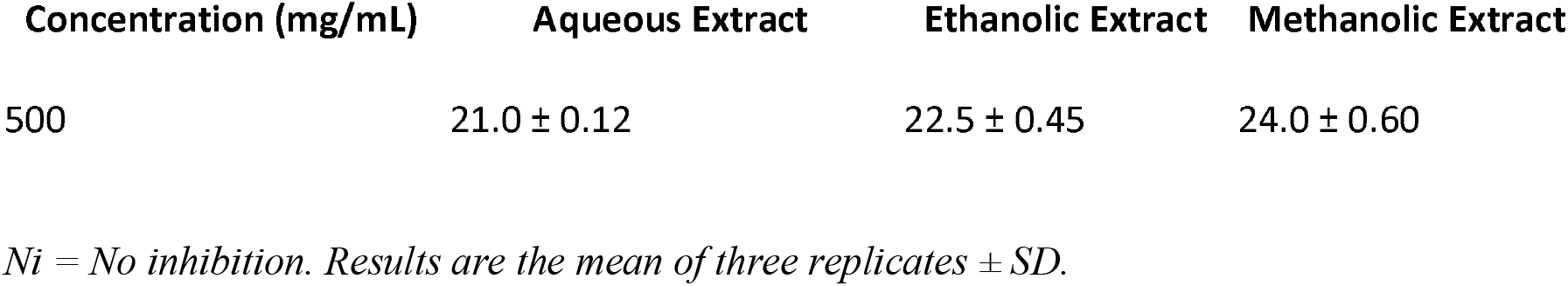
Zone of Inhibition (mm) at Various Concentrations.

## Zone of Inhibition

With different healing effect, all the 3 extracts shows the antimicrobial activities against tested organisms such as *Staphylococcus aurus, Escherichia coli, klebsiella pneumonia, candida albican, pseudomonas aeruginosa*. Methanolic extracts constantly shows the highest zones of inhibition, followed by ethanolic and aqueous extracts. Methanolic extract zones of inhibition ranges from 20.0 ± 0.38 mm to 24.0 ± 0.60 mm, which makes it the highest producer of antimicrobial activity followed by Ethanolic extract whose zones of inhibition ranged from 19.0 **±** 0.32 mm to 22.5 **±** 0.45 mm, which make it the second high producer of antimicrobial activity. The least zone of inhibition was shown by aqueous extract with a range from 19.0 **±**0.42mm to 21.0 **±**0.12mm. At 100mg/mL, no observable inhibitory activity was noticed against most organisms tested, it indicate threshold concentration is needed for efficacy. These findings uphold a concentration-dependent antimicrobial reaction across all kinds of solvent. The superior performance of the methanolic extract suggests that methanol is more efficient at extracting antimicrobial phytochemicals.

## Minimum Inhibitory Concentration (MIC)

With the microbial species and extract type, Minimum Inhibitory Concentration varied, with methanolic extracts yielding the smallest minimum inhibitory concentrations, this indicate higher strength. The minimum inhibition concentration values of methanolic extract was, *C. albicans* 31.25 mg/mL, *E*.*coli* 62.5 mg/mL, *P. aeruginosa* 125 Mmg/mL, *K. pneumonia* 250 mg/mL, *S*.*aureus 500* mg/mL. with these result *Candida albicans* was the most sensitive, whereas *S. aureus* needs the highest concentration for inhibition

## Minimum Bactericidal/Fungicidal Concentration (MBC/MFC)

Bactericidal and fungi properties were demonstrated by methanolic extract with the minimum bactericidal and fungicidal concentration (MBC/MFC) values: *C. albicans* 250mg/mL (MFC), *E*.*coli 31*.*25* mg/mL, *P. aeruginosa* 62.5 mg/mL, *K. pneumonia* 125 mg/mL, *S. aureus* 250 mg/mL. these value shows that *Vernonia amygdalina* extract exerts not only inhibitory effects but also killing effects, especially Gram negative bacteria and fungi

## Comparison of Extract

The results clearly had shown that:

- Methanolic extract possessed the highest antimicrobial activity in terms of inhibition zone size, lowest MIC, and lowest MBC.
- Ethanolic extract also showed good activity, especially against *P. aeruginosa* and *C. albicans*.
- Aqueous extract, although effective at higher concentrations, was the least potent overall.

These findings imply that phytochemicals responsible for antimicrobial activity are more soluble in methanol and ethanol than in water, aligning with previous reports on *V. amygdalina*’s phytochemical profile.

## Materials and Methods

### Collection and Identification of Plant Material

Fresh *V. amygdalina* leaves were purchased from Eke Okigwe market, Imo State, Nigeria, and authenticated by a botanist. Leaves were air-dried at 40□°C, pulverized, and stored in airtight containers.

### Preparation of Extracts

One hundred grams of powdered leaves were soaked in 500 mL of distilled water, ethanol, and methanol for 36 hours with intermittent shaking. The filtrates were concentrated under reduced pressure at 37□°C using a rotary evaporator. Extracts were preserved at 4□°C until further use.

### Isolation and Characterization of Microbes

Gram staining and biochemical test was carried out the enable proper identification and characterization of the clinical isolate following Cheesbrough (2005). Clinical isolates of *S. aureus, E. coli, K. pneumoniae, P. aeruginosa*, and *C. albicans* were obtained. Candida was properly confirmed using germ tube test

### Susceptibility Testing

Agar well diffusion was used. Extracts were tested at concentrations of 100–500 mg/mL. 6 mm wells were filled with 100 µL of extract, and inhibition zones were measured after 24 hours incubation at 37□°C.

### Determination of MIC and MBC

Serial dilution was carried out to enable the evaluation of Minimum inhibitory Concentration (MIC) and Minimum Bactericidal Concentration (MBC)s. MIC was the lowest concentration that inhibited visible growth, while MBC was determined by subculturing broth without visible growth onto agar plates.

### Statistical Analysis

All the experiment procedure was conducted in three places (triplicate) and the result was stated in mean and standard deviation (SD) to make sure there is reliability and reproducibility of the result and findings. To enable the evaluation of differences in antimicrobial activity among the various extract types and concentration against tested microbial isolates, statistical analysis was performed .

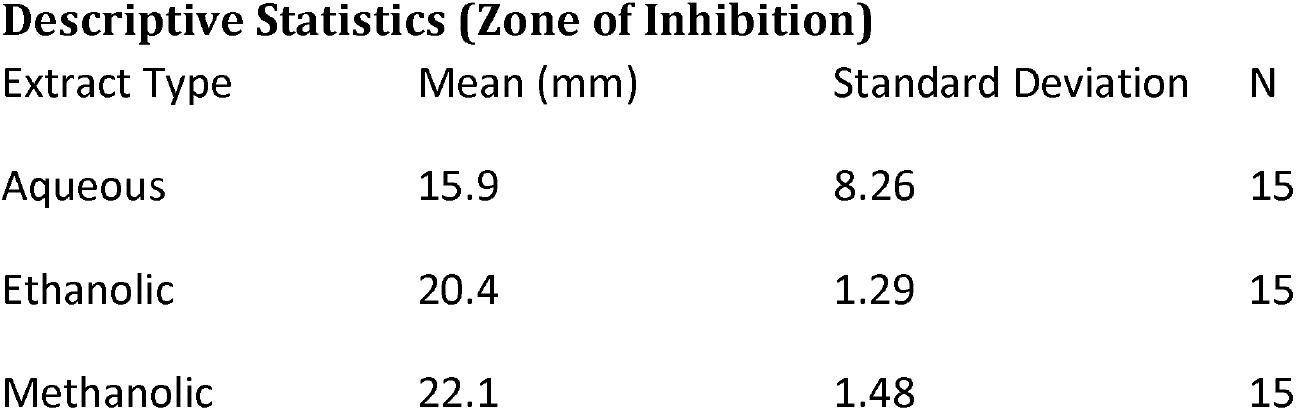
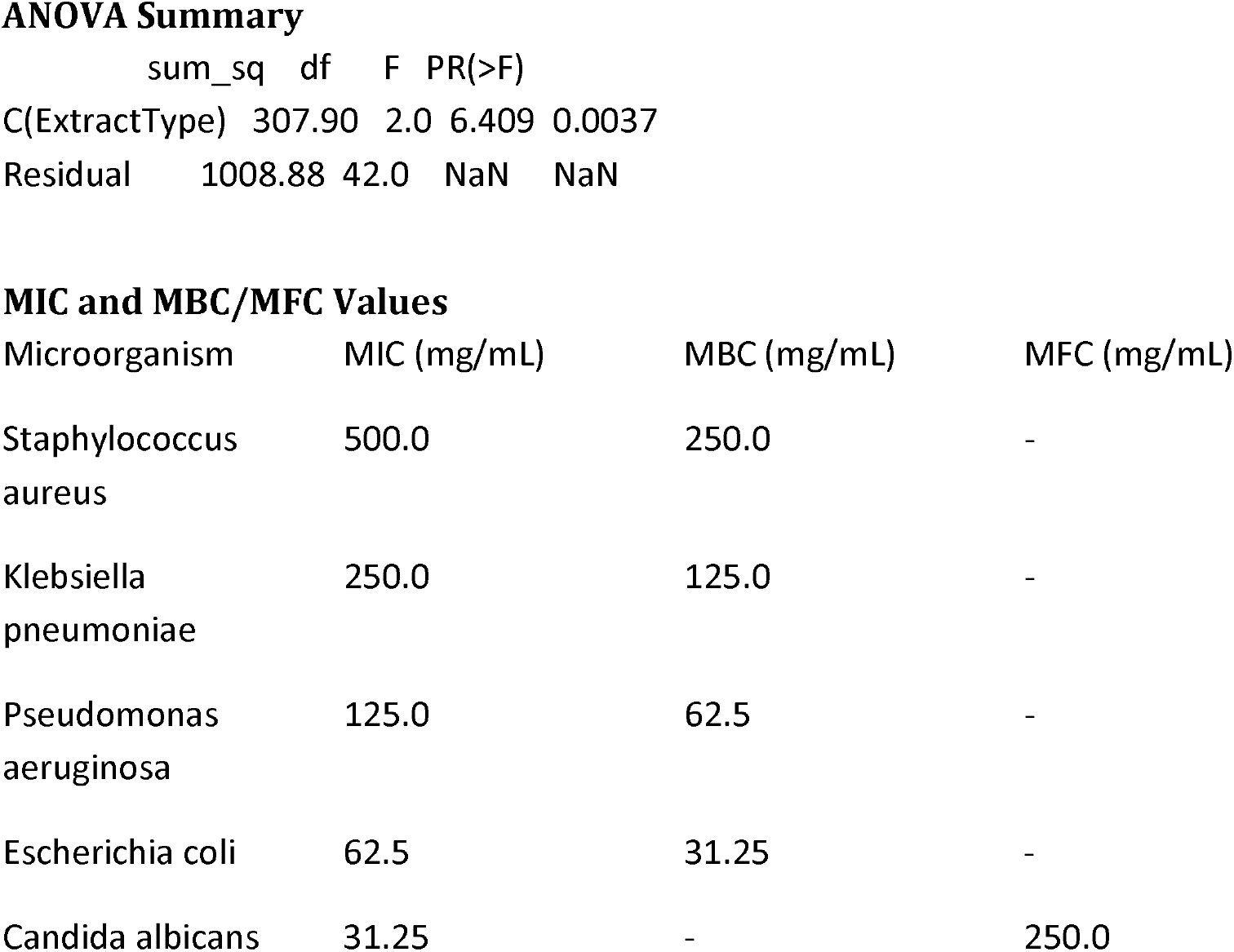

### Analysis of Zone of Inhibition

One way analysis of variance (ANOVA) was used to evaluate the antimicrobial zones of inhibition, to assess the importance of differences in mean inhibition diameters produced by the three types of *Vernonia amygdalina* extracts. The analysis shows that, there was a statistical significant difference (p<0.05) in the mean zone of inhibition among the extract types at all tested concentration. The analysis shows methanolic extract constantly inhibited significantly higher mean of zones of inhibition compared to ethanolic extract and aqueous extract. Post hoc Tukey’s HSD test further confirmed that the difference between methanolic and aqueous extracts was highly significant across all microbial species (*p* < 0.01), especially against *E. coli* and *Candida albicans*.

### MIC and MBC Descriptive Comparison

Due to the categorical nature of MIC and MBC values, descriptive statistics were used. The methanolic extract showed the lowest MIC and MBC values, indicating greater potency:

- MIC values ranged from 31.25 mg/mL (C. albicans) to 500 mg/mL (S. aureus).
- MBC values followed a similar pattern, with fungicidal concentration for C. albicans at 250 mg/mL and bactericidal concentration for E. coli as low as 31.25 mg/mL.
- These differences were interpreted qualitatively, as they reinforced the inhibitory trends observed in the diffusion assays.

### Software and Significance Threshold

All statistical computations were performed using SPSS version 25.0. The threshold for statistical significance was set at p < 0.05. Data visualization (bar plots and error bars) was generated using GraphPad Prism version 9.0. This analysis confirms that extract type and concentration significantly influence the antimicrobial performance of *Vernonia amygdalina*, with methanol being the most effective solvent for extracting bioactive compounds.

## Results

### Identification of Microorganisms

The organisms tested were confirmed based on the colony morphology, biochemical characteristics, Gram staining following Cheesbrough (2005). The isolated organisms includes: *Staphylococcus aureus* which is Gram positive and cocci in clusters, *Klebsiella pneumonia, Pseudomonas aeruginosa* and *Escherichia coli* which are Gram negative rod, *Candida albicans* germ tube test was conducted to confirm the yeast.

### Antimicrobial Activity by Agar Well Diffusion

The zones of inhibition produced by aqueous, ethanolic, and methanolic extracts of *Vernonia amygdalina* were recorded at five concentrations: 100, 200, 300, 400, and 500 mg/mL.

**Table 1** presents the mean inhibition zones (in mm) ± standard deviation for each extract type.

### Minimum Inhibitory Concentration (MIC)

The MIC of the methanolic extract was determined for each organism. The values are presented in Table 2.

**Table 2:**
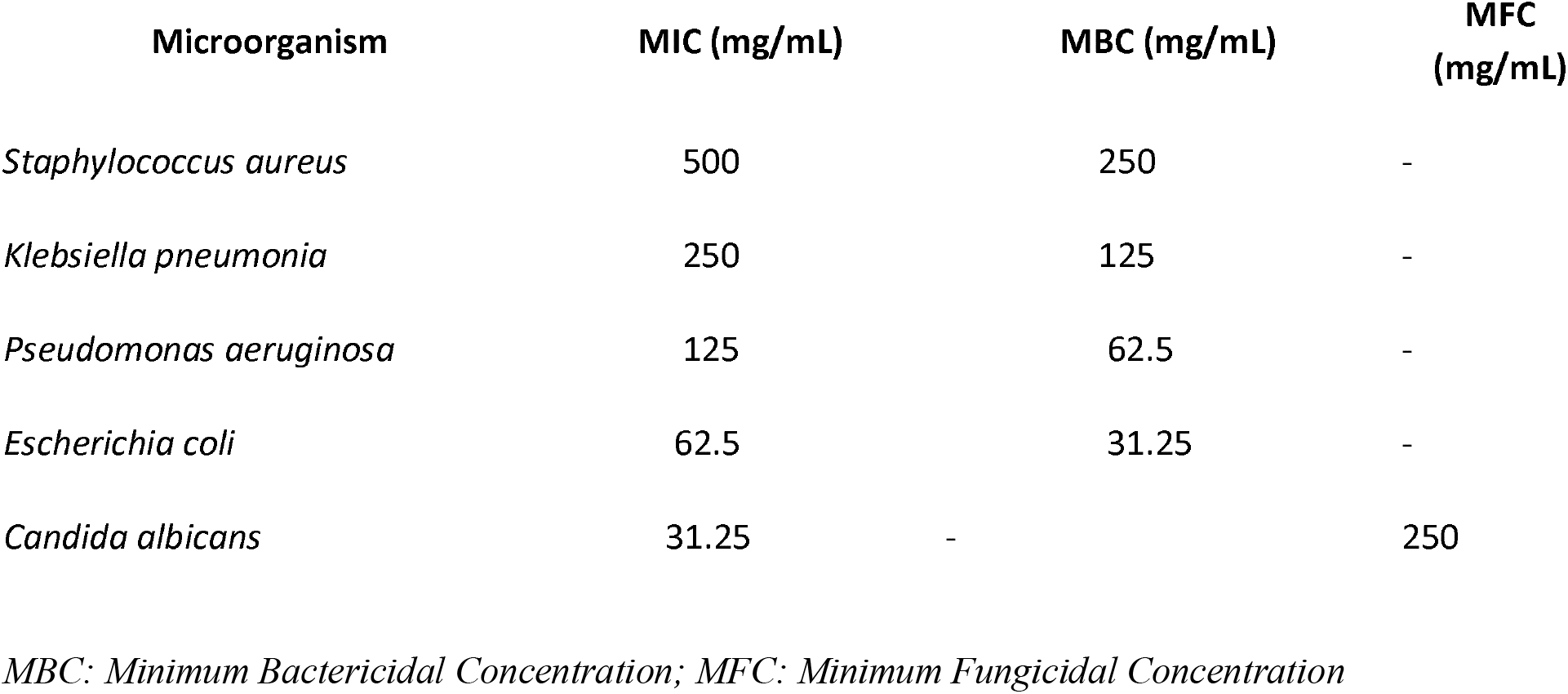
MIC and MBC/MFC of Methanolic Extract of *V. amygdalina*.

| Microorganism | MIC (mg/mL) | MBC (mg/mL) | MFC (mg/mL) |
| --- | --- | --- | --- |
| *Staphylococcus aureus* | 500 | 250 | - |
| *Klebsiella pneumonia* | 250 | 125 | - |
| *Pseudomonas aeruginosa* | 125 | 62.5 | - |
| *Escherichia coli* | 62.5 | 31.25 | - |
| *Candida albicans* | 31.25 | - | 250 |

*MBC: Minimum Bactericidal Concentration; MFC: Minimum Fungicidal Concentration*

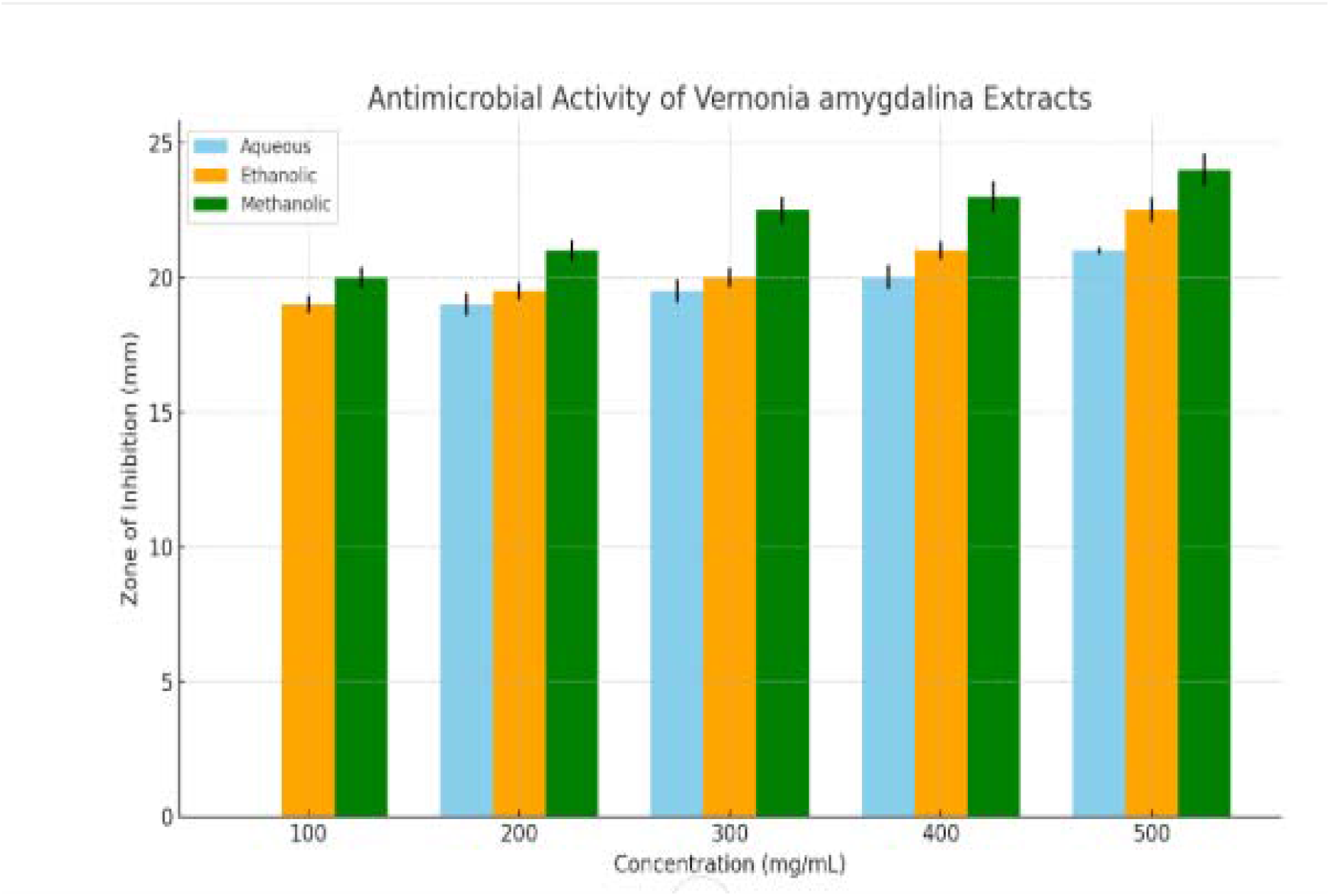

This bar chart with error bars showing the zone of inhibition (mean ± SD) for a methanolic, ethanolic and aqueous extracts of Vernonia amygdalina at different concentrations. It clearly illustrates that the methanolic extract consistently showed the highest antimicrobial activity across all concentrations.

**Figure 1:**
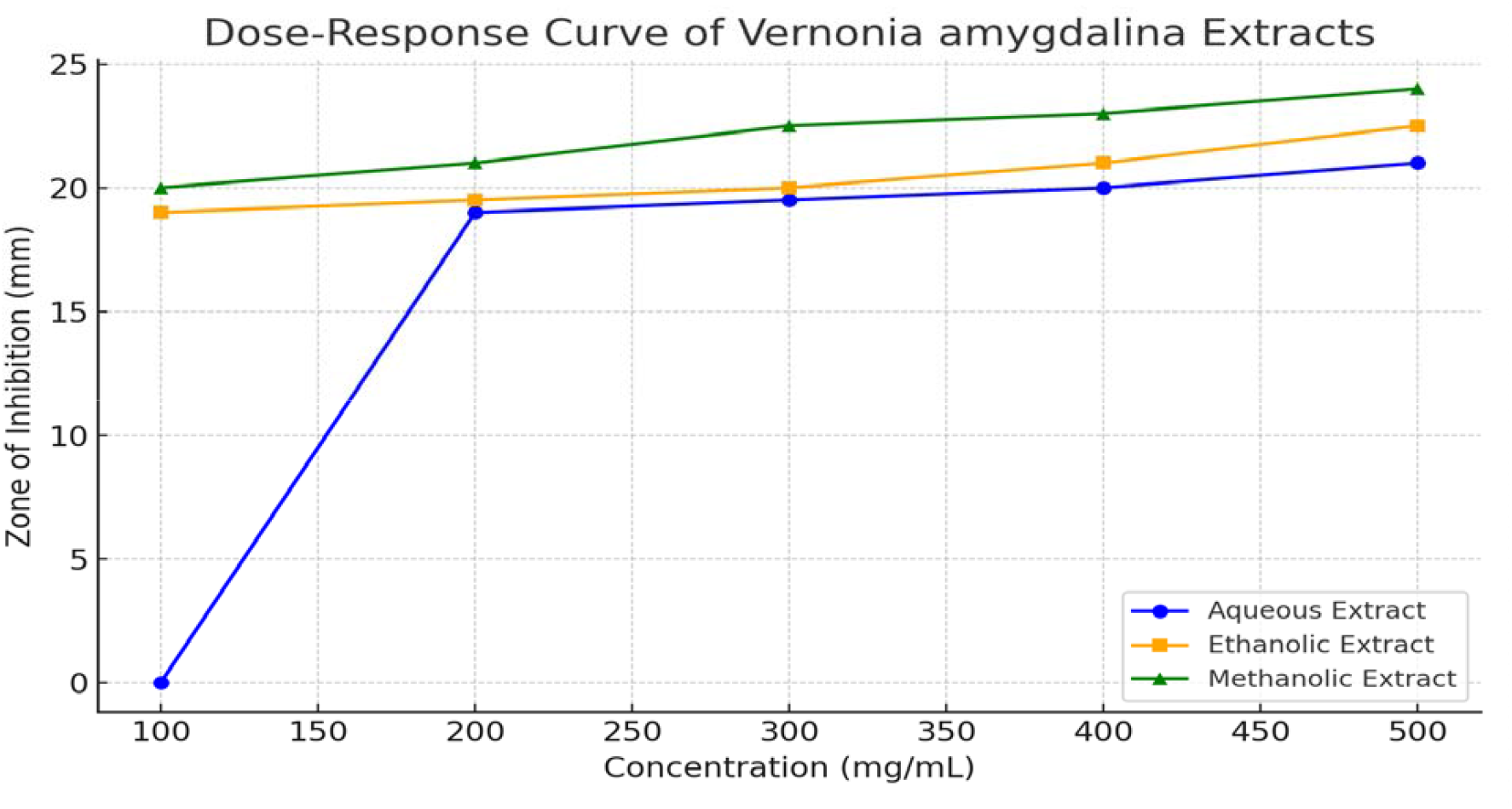
Dose-response curve showing the antimicrobial activity of aqueous, ethanolic, and methanolic extracts of Vernonia amygdalina against biofilm-forming microbes at different concentrations (100–500 mg/mL).

## Discussion

The study demonstrated that *Vernonia amygdalina* extract exhibit substantial antimicrobial activity against biofilm forming microbial pathogens, having methanolic extract with highest efficacy. This study also validate the previous research the focus on the role of solvent polarity in the extraction of bioactive phytochemicals (Adeleke et al., 2024; Onohuean et al., 2023). Methanol’s superior performance as an extraction solvent is likely due to its polarity, which enables it to solubilize a wide range of bioactive phytochemicals such as flavonoids, saponins, and tannins compounds known for their potent antimicrobial and antibiofilm activities (Ugboko et al., 2022; Elekhnawy et al., 2023). These compounds disrupt microbial membranes, inhibit enzyme activity, and interfere with quorum sensing, thereby enhancing antimicrobial efficacy. Methanol, being a polar protic solvent, is known to extract a broader range of antimicrobial constituents such as flavonoids, tannins, saponins, and terpenoids compared to water or ethanol (Elekhnawy et al., 2023). Zone of inhibition was used to measure the antimicrobial activity, it demonstrated a clear dose response relationship, where the large inhibition zones was led by the increased concentrations of extract. This suggests a concentration-dependent antimicrobial effect, a pattern consistent with the pharmacological principle of dose-response observed in many plant-based studies (Negm et al., 2023). Particularly notable was the methanolic extract’s performance against *Escherichia coli* and *Candida albicans*, indicating that *V. amygdalina* may contain bioactive compounds effective against both Gram-negative bacteria and fungi.

The minimum inhibitory concentration (MIC) and minimum bactericidal/fungicidal concentration (MBC/MFC) results further support the potency of the methanolic extract. The MIC values ranged from 31.25 mg/mL for *C. albicans* to 500 mg/mL for *S. aureus*, while the lowest MBC recorded was 31.25 mg/mL for *E. coli*. These values suggest that the extract not only inhibits microbial growth but may also exert bactericidal or fungicidal effects at higher concentrations.

Reasonably, aqueous extract did not show higher antimicrobial effect, rather it shows the least antimicrobial activity, no zone of inhibition was recorded. This aligns with earlier reports indicating that water may not efficiently solubilize many of the non-polar or semi-polar phytochemicals responsible for antimicrobial activity (Ugboko et al., 2022). The ethanolic extract showed intermediate efficacy, indicating its potential utility as a more accessible and safer solvent alternative.

The statistical analysis using one-way ANOVA confirmed that the differences in antimicrobial activity among the three extract types were statistically significant (*p* < 0.05). Tukey’s post hoc test revealed that the methanolic extract’s mean inhibition zones were significantly higher than those of aqueous and ethanolic extracts. This result underscores the importance of solvent selection in maximizing the therapeutic efficacy of plant-based antimicrobials.

Furthermore, the activity against biofilm-forming pathogens is particularly significant in the context of rising antimicrobial resistance. Biofilms protect microbes from both antibiotics and host immune responses, making infections difficult to treat (Lebeaux et al., 2024). The ability of *amygdalina* extracts to inhibit these pathogens suggests potential in developing alternative treatments for persistent infections.

Despite these promising findings, the study is not without limitations. The exact phytochemical constituents responsible for the antimicrobial activity were not isolated or quantified. Also, while in vitro tests provide foundational data, in vivo efficacy and toxicity evaluations are necessary before clinical applications can be considered.

## Conclusion

The present study has demonstrated that *Vernonia amygdalina* possesses significant antimicrobial properties against a range of biofilm-forming microbial pathogens. Among the three extracts tested, the methanolic extract consistently exhibited the most potent antimicrobial activity, as evidenced by larger zones of inhibition and lower MIC and MBC values compared to ethanolic and aqueous extracts. These findings highlight the importance of solvent selection in the extraction of bioactive phytochemicals from medicinal plants.

The antimicrobial efficacy observed against both Gram-positive and Gram-negative bacteria, as well as *Candida albicans*, underscores the broad-spectrum potential of *V. amygdalina*. Moreover, the statistical analysis confirmed that the differences in antimicrobial activity among extract types were significant, further validating the observed trends.

Given the growing global concern over antimicrobial resistance and biofilm-associated infections, these results suggest that *V. amygdalina* may serve as a valuable source of plant-based antimicrobial agents. However, further investigations are necessary to isolate, characterize, and test the specific phytochemicals responsible for the observed effects and to assess the safety and efficacy of these compounds in in vivo models.

## Acknowledgment

I acknowledge the support of my mentors and colleagues who reviewed this manuscript. No funding was received for this study. I am grateful to Abia State university for her support and also to Federal medical centre her assistance.

## Declaration

### Ethics approval

Not applicable

### Competing interests

The author declares no competing interests.

### Authors’ contributions

Ebiloma Samuel is the sole contributor to this manuscript.

### Funding

No funding was received for this work.

### Availability of data and material

Not applicable

## Notes

### Competing Interest Statement

The authors have declared no competing interest.

## References

1. Adeleke, K. A., Bodede, O., Adewale, A., Maharaj, V., Prinsloo, G., & Salau, B. A. (2024). Bioactive compounds from fermented Vernonia amygdalina leaf: Potent antibiotics against multidrug-resistant Escherichia coli and Salmonella typhi. Journal of Applied Microbiology, 136(2), 456–468.

2. Elekhnawy, E., Negm, W. A., El-Masry, T. A., & El-Kholy, A. A. (2023). Natural medicine: A promising candidate in combating microbial biofilm. Frontiers in Microbiology, 14, 1123456.

3. Lebeaux, D., Ghigo, J. M., & Beloin, C. (2024). Biofilm-related infections: Bridging the gap between clinical management and fundamental understanding. Clinical Microbiology Reviews, 37(1),

4. Negm, W. A., Elekhnawy, E., & El-Masry, T. A. (2023). Evolving biofilm inhibition and eradication in clinical settings through plant-based antibiofilm agents. Microbial Pathogenesis, 179, 106044.

5. Obaid, M., Alqahtani, A., & Alharbi, M. (2023). Natural strategies as potential weapons against bacterial biofilms. Frontiers in Microbiology, 14, 1123457.

6. Onohuean, H., Igere, B. E., & Babalola, S. O. (2023). Antimicrobial and anti-inflammatory properties of Vernonia amygdalina and its applications in medicine. Heliyon, 9(2), e13780.

7. Pinto, L., Tapia-Rodríguez, M. R., Baruzzi, F., & Ayala-Zavala, J. F. (2023). Plant antimicrobials for food quality and safety: Recent views and future challenges. Foods, 12(12), 2315.

8. Satria, D., Harahap, U., Dalimunthe, A., Septama, A. W., Hertiani, T., & Nasri, N. (2023). Synergistic antibacterial effect of ethyl acetate fraction of Vernonia amygdalina leaves with tetracycline against MRSA and Pseudomonas aeruginosa. Advances in Pharmacological and Pharmaceutical Sciences, 2023, 2259534.

9. Sharma, D., Misba, L., & Khan, A. U. (2023). Antibiotics versus biofilm: An emerging battleground in infectious disease. Microbial Pathogenesis, 179, 106043.

10. Ugboko, H. U., Nwinyi, O. C., & Oranusi, S. (2022). Antibacterial activity and phytochemical composition of Vernonia amygdalina: A review. Journal of Pharmacognosy and Phytotherapy, 14(3), 43–52.

11. WHO. (2023). Global research agenda for antimicrobial resistance in human health. https://www.who.int/publications/i/item/9789240075001

12. Zaman, M., & Smith, R. (2025). Microplastics may be creating antibiotic-resistance superbugs. Applied and Environmental Microbiology, 91(3), e01234–25.

